# Mechanical strengthening of cell-cell adhesion during mouse embryo compaction

**DOI:** 10.1101/2023.12.07.570568

**Authors:** Ludmilla de Plater, Julie Firmin, Jean-Léon Maître

## Abstract

Compaction is the first morphogenetic movement of the eutherian mammals and involves a developmentally regulated adhesion process. Previous studies investigated cellular and mechanical aspects of compaction. During mouse and human compaction, cells spread onto each other as a result of a contractility-mediated increase in surface tension pulling at the edges of their cell-cell contacts. However, how compaction may affect the mechanical stability of cell-cell contacts remains unknown. Here, we used a dual pipette aspiration assay on cell doublets to quantitatively analyze the mechanical stability of compacting mouse embryos. We measured increased mechanical stability of contacts with rupture forces growing from 40 to 70 nN, which was highly correlated with cell-cell contact expansion. Analyzing the dynamic molecular reorganization of cell-cell contacts, we find minimal recruitment of the cell-cell adhesion molecule Cdh1 (also known as E-cadherin) to contacts but we observe its reorganization into a peripheral adhesive ring. However, this reorganization is not associated with increased effective bond density, contrary to previous reports in other adhesive systems. Using genetics, we reduce the levels of Cdh1 or replace it with a chimeric adhesion molecule composed of the extracellular domain of Cdh1 and the intracellular domain of Cdh2 (also known as N-cadherin). We find that reducing the levels of Cdh1 impairs the mechanical stability of cell-cell contacts due to reduced contact growth, which nevertheless show higher effective bond density than WT contacts of similar size. On the other hand, chimeric adhesion molecules cannot form large or strong contacts indicating that the intracellular domain of Cdh2 is unable to reorganize contacts and/or is mechanically weaker than the one of Cdh1 in mouse embryos. Together, we find that mouse embryo compaction mechanically strengthens cell-cell adhesion via the expansion of Cdh1 adhesive rings that maintain pre-compaction levels of effective bond density.

## Introduction

During preimplantation development, mammalian embryos form the blastocyst, a structure sharing many conserved features among mammals that, in eutherians and some marsupials, is required for implantation (1–4). The eutherian blastocyst forms after a series of morphogenetic movements, the first one being compaction. Compaction is a developmentally regulated adhesion process during which cells of the mammalian embryo come into close contact (5). Before compaction, cells of mammalian embryos appear loosely attached to one another and cell-cell contacts do not grow substantially between cleavage divisions. In the mouse, compaction takes place at the 8-cell stage within 8-10 h of the 3^rd^ round of cleavage divisions (6). Some of the cellular and mechanical mechanisms underlying compaction have been characterized. Compaction requires the cell-cell adhesion molecule Cdh1, also known as E-cadherin and initially called *uvomorulin* after it was discovered in compacting mouse embryos (7). After this striking discovery, changes in Cdh1-mediated cell-cell adhesion were proposed to drive compaction and multiple components of adherens junctions were investigated during compaction (8–11). Changes in adhesion molecules were thought to drive compaction by increasing adhesion energy, in a way similar to what takes place in contacting soap bubbles (12). Assuming cells of compacting embryos behave as liquid droplets, increasing adhesion energy reduces the tension at cell-cell contacts γ_cc_and lowers the ratio of tensions at the cell-cell and cell-medium interfaces γ_cc_/ γ_cm_(13, 14). Following the Young-Dupré equation cos (θ/2) = γ_cc_/ 2γ_cm_, this increases the external angle of contact θ and drives compaction (15). However, estimations of the adhesion energy provided by cadherin adhesion molecules, such as Cdh1, were found to be too weak when compared to another powerful morphogenetic engine in animal cells: actomyosin contractility (16, 17). In mouse and human embryos, actomyosin contractility is also required for compaction (6, 18, 19). Actomyosin contractility raises the surface tension at the cell-medium interface γ_cm_, which changes the ratio of surface tensions between the cell-cell and cell-medium interfaces γ_cc_/ γ_cm_(6, 19). This increases the contact angle θ between cells and compacts the embryo. In maternal Cdh1 mutants (20), which lack Cdh1 at the time of compaction, cells do not expand their cell-cell contacts (6). Despite the absence of Cdh1, contractility increases the surface tension at the cell-medium interface γ_cm_. However, contrary to wildtype embryos, contractile forces cannot be transmitted between cells without Cdh1. Therefore, compaction results from increased contractile forces that pull cells together via Cdh1 mechanical anchors (5).

During compaction, mechanical coupling between cells is expected to strengthen. Indeed, cell-cell contacts become larger and should lead to an increased number of bonds connecting cells, provided that bond density remains at least at the same level. In fact, when forming a cell-cell contact, cadherin adhesion molecules become enriched and bond density often increases (21). In some cases, cadherin adhesion molecules cluster at the edges of the cell-cell contact, which, in doublet of cells, forms a ring of adhesion molecules where it can engage with the cytoskeleton and modulates the effective bond density (16, 22). Therefore, the relationship between cell-cell contact area and the number of adhesion molecules holding cells together may not be trivial. Adhesion rings are thought to act as a scaffold on which actomyosin contractility can effectively pull and extend the cell-cell contact area. Pulling on adhesion complexes at the edges of cell-cell contacts can also lead to mechanosensitive reinforcement of adherens junctions. Mechanosensitive strengthening can occur via the recruitment of adhesion molecules (22–24) and/or via their anchoring to the cytoskeleton (25, 26). Cadherin adhesion molecules are connected to the actin cytoskeleton via several proteins including α-catenin, which can unfold when stretched (25, 27). This leads to increased connections with the actin cytoskeleton and mechanically strengthens cell-cell contacts by changing the effective bond density. Therefore, during compaction, cell-cell contacts could be mechanically stabilized by increased contact surface, increased adhesion molecule recruitment, adhesion molecule reorganization at the contact rims, and/or increased coupling of individual adhesion complexes. Each of these parameters are susceptible to modulate the effective bond density and to determine the mechanical stability of cell-cell contacts. Taken together, while the mechanics of cell-cell contact spreading are well studied, how mechanical coupling may change during compaction is not.

Assessing the mechanical coupling of cell-cell adhesion can be done by separating cells while measuring the force necessary to do so (14, 28). This is typically performed in doublets of cells forming a single cell-cell contact. Assays based on atomic force microscopes (AFM) have measured adhesive coupling between cells up to tens of nN, which are typically reached within a few minutes of contact (29, 30). While AFM allows for highly accurate measurements, stronger cell-cell contacts can be difficult to separate using AFM (14, 28). To measure adhesive coupling up to hundreds of nN, dual pipette aspiration (DPA) assays can be used (31, 32). Also, DPA is easily compatible with light microscopy, which allows one to obtain information about how contacts reorganize during separation. Notably, separation of cell doublets often leads to the formation of membrane tethers connecting separated cells (16, 24). This indicates that some adhesion complexes remain attached in trans and rather detach intracellularly. The weakest link of the adhesion complex may be located at the level of α-catenin or of the actin cytoskeleton, depending on the cellular context (16, 24). In fact, modifying the intracellular binding of cadherins to the cytoskeleton directly affects mechanical coupling (16, 32). This can also be measured when comparing the intracellular domains of different cadherin adhesion molecules, with Cdh1 intracellular domain showing stronger coupling than that of Cdh7 as shown, for example, using chimeric adhesion molecules (33). Interestingly, in mouse embryos, replacing Cdh1 by a chimeric adhesion molecule, called EcNc, composed of the extracellular domain of Cdh1 and the intracellular domain of Cdh2, also known as N-cadherin, affects preimplantation development (34). However, the precise effect on compaction and in particular on the mechanical coupling of cells expressing an EcNc chimeric adhesion molecule has not been characterized.

Finally, another function of adhesion molecules is to signal to reorganize the underlying cytoskeleton (35). This can have multiple effects on contacting cells such as contact inhibition of locomotion in migratory cells (36, 37). In particular, cadherin signaling via small GTPases leads to the downregulation of actomyosin contractility at the cellular level (38) or locally at cell-cell contacts (39, 40). During compaction, this is key to reduce or maintain contact contractility and associated tension γ_cc_ at low levels, as seen in mouse or human embryos with impaired Cdh1-based adhesion (6, 19). The ability of different cadherin adhesion molecules to signal to the actomyosin cytoskeleton is mostly unknown and it is for example unclear how an EcNc chimeric adhesion molecule would affect contact expansion during compaction as compared to Cdh1.

Here, we set out to study the mechanical coupling of cells during mouse embryo compaction. Using a DPA assay, we measured the mechanical coupling of compacting blastomeres and analyzed its relationship to contact size, adhesion molecule recruitment and reorganization. We find that mechanical coupling increases during compaction and can be almost entirely explained by contact growth. We further measure that adhesion molecules do not increase in concentration at the growing cell-cell contacts but become enriched in a ring located at their periphery. This reorganization of adhesion molecule is not associated with increased mechanical coupling. Instead, using mice expressing lower levels of Cdh1 or an EcNc chimeric adhesion molecules knocked in the Cdh1 locus, we find that both contact expansion and mechanical coupling are impacted by the intracellular binding of adhesion molecules. Together, this study of compaction provides valuable insights into the regulation of mechanical coupling during physiological increase in cell-cell adhesion.

## Material and methods

### Embryo work

#### Recovery and culture

All animal work is performed in the animal facility at the Institut Curie, with permission by the institutional veterinarian overseeing the operation (APAFIS #11054-2017082914226001 and APAFIS #39490-2022111819233999 v2). The animal facilities are operated according to international animal welfare rules.

Embryos are isolated from superovulated female mice mated with male mice. Superovulation of female mice is induced by intraperitoneal injection of 5 international units (IU) pregnant mare’s serum gonadotropin (Ceva, Syncro-part), followed by intraperitoneal injection of 5 IU human chorionic gonadotropin (MSD Animal Health, Chorulon) 44-48 hours later. Embryos are recovered at E1.5 by flushing oviducts with 37°C FHM (Millipore, MR-122-D) using a modified syringe (Acufirm, 1400 LL 23).

Embryos are handled using an aspirator tube (Sigma, A5177-5EA) equipped with a glass pipette pulled from glass micropipettes (Blaubrand intraMark or Warner Instruments).

Embryos are placed in KSOM (Millipore, MR-107-D) or FHM supplemented with 0.1 % BSA (Sigma, A3311) in 10 μL droplets covered in mineral oil (Acros Organics). Embryos are cultured in an incubator with a humidified atmosphere supplemented with 5% CO2 at 37°C. To remove the Zona Pellucida (ZP), embryos are incubated for 45-60 s in pronase (Sigma, P8811) at the 2-or 4-cell stage.

To dissociate blastomeres, 4-cell stage embryos without ZP are aspirated 1-3 times into a glass pipette of size larger than a 4-cell stage blastomere and smaller than the whole embryo (i.e. ∼50-80 μm in diameter).

For imaging, embryos, dissociated 4-cell stage blastomeres or 8-cell stage doublets are placed in 3.5 or 5 cm glass-bottom dishes (MatTek).

#### Mouse lines

Mice are used from 5 weeks old on. (C57BL/6xC3H) F1 hybrid strain is used for wild type (WT).

To remove LoxP sites specifically in oocytes, Zp3-cre (Tg(Zp3-cre)93Knw) mice are used (41). To generate mCdh1^+/-^ embryos, Cdh1^tm2kem^mice are used (42) to breed Cdh1^tm2kem/tm2kem^; Zp3^Cre/+^mothers with WT fathers.

To generate mCdh1^EcNc/-^ embryos, Cdh1^tm2kem^mice are used (42) to breed Cdh1^tm2kem/ tm2kem^; Zp3^Cre/+^mothers with Cdh1^tm3.1Mpst/+^fathers (34). To genotype EcNc embryos, we used antibodies recognizing the extra- or intra-cellular domains of Cdh1, or the HA tag added to the EcNc (34).

#### Immunostaining

Embryos or doublets are fixed in 2% PFA (Euromedex, 2000-C) for 10 min at 37°C, washed in PBS and permeabilized in 0.01% Triton X-100 (Euromedex, T8787) in PBS (PBT) at room temperature before being placed in blocking solution (PBT with 3% BSA) at 4°C for 2-4 h. Primary antibodies are applied in blocking solution at 4°C overnight. After washes in PBT at room temperature, embryos are incubated with secondary antibodies, DAPI and/or phalloidin in blocking solution at room temperature for 1 h. Embryos are washed in PBT and imaged immediately after.

| Primary antibody | Dilution | Provider | RRID |
| --- | --- | --- | --- |
| Cdh1 (DECMA-1) Rat monoclonal | 1:200 | eBioscience, 14-3249-82 | AB_1210458 |
| Cdh1 (24E10) Rabbit monoclonal | 1:200 | Cell Signaling 3195 | AB_2291471 |
| Anti-HA.11 Tag mouse monoclonal | 1:200 | Biolegend 901513 | AB_2565336 |

| Secondary antibodies and dyes | Dilution | Provider | RRID |
| --- | --- | --- | --- |
| Alexa Fluor Plus 488 Goat anti-mouse | 1:200 | Invitrogen, A32723 | AB_2633275 |
| Alexa Fluor 546 Goat anti-mouse | 1:200 | Invitrogen, A11003 | AB_2534071 |
| Alexa Fluor Plus 488 Goat anti-rabbit | 1:200 | Invitrogen, A32731 | AB_2633280 |
| Alexa Fluor Plus 546 Goat anti-rabbit | 1:200 | Invitrogen, A11010 | AB_2534077 |
| Alexa Fluor 546 Goat anti-rat | 1:200 | Invitrogen, A11081 | AB_141738 |
| Alexa Fluor 633 phalloidin | 1:200 | Invitrogen, A22284 |  |
| 4',6-diamidino-2-phenylindole (DAPI) | 1:1000 | Invitrogen, D1306 | AB_2629482 |

### Microscopy

Time lapse imaging of dissociated 4-cell stage blastomeres and separation of 8-cell stage doublets are performed on a Leica DMI6000 B inverted microscope equipped with a 40x/0.8 DRY HC PL APO Ph2 (11506383) objective. A 0.7x lens is mounted in front of a Retina R3 camera. The microscope is equipped with a custom incubation chamber (EMBLEM) to keep the sample at 37°C and maintain the atmosphere at 5% CO_2_.

Time lapse imaging of dissociated 4-cell stage blastomeres without separation at the 8-cell stage are performed on a Celldiscoverer 7 (Zeiss) equipped with a 20x/0.95 objective. A 2x tubelens in front of an Axiovert (Zeiss) camera. The microscope is equipped with an incubation chamber keeping the sample at 37°C and maintaining the atmosphere at 5% CO_2_.

Immunostainings are imaged on an inverted Zeiss Observer Z1 microscope with a CSU-X1 spinning disc unit (Yokogawa). Excitation is achieved using 405, 488, 561 and 642 nm laser lines through a 63x/1.2 C Apo Korr water immersion objective; emission is collected through 450/50 nm, 525/50 nm, 595/50 band pass or 610 nm low pass filters.

### Micropipette aspiration

#### Micropipettes preparation

To forge the micropipettes, glass capillaries (World Precision Instruments TW100-3) are pulled using a P-97 Flaming Brown needle puller (Sutter Instrument) with the following settings: Ramp +5, Pull 50, Velocity 50, Time 50 and Pressure 500.

Using a microforge (Narishige, MF-900), needles are cut to form a blunt opening of radius 6-9 μm and bent 80-100 μm away from the tip at a 20° angle.

#### Dual pipette aspiration setup

Micropipettes are mounted on a micromanipulator (Leica AM6000) using a grip head and capillary holder (Eppendorf, 920007392 and 9200077414). The micropipettes are connected to PBS-filled intermediate reservoirs of which the height are controlled using a 50 mm microscale translating stage (Newport) to generate positive and negative pressures (43). The intermediate reservoirs are connected to a microfluidic pump (Fluigent, MFCS-EZ) delivering negative pressures with a 2.5 Pa resolution. The pressure is controlled using Maesflow software (Fluigent). The output pressure is calibrated by finding the height of the intermediate reservoir at which no flow in the micropipette is observed (using floating particles found in the dish, ‘no flow’ is considered achieved when the position of the particle inside the micropipette is stable for ∼10 s and, if slow drift can be reverted with 10 Pa).

The separation force *F*_*s*_ is determined as previously described (31, 43). A doublet is grabbed by two glass micropipettes (holding and probing pipettes) on the opposite sides of the cell-cell contact. The holding micropipette is used to firmly hold one cell with a fixed pressure ranging from 100-550 Pa. The probing micropipette is used to apply stepwise increasing pressures ranging from 10-1000 Pa with step-ranges between 10 and 100 Pa to the other cell. After each pressure step, micropipettes are pulled apart at 60 μm/s using MetaMorph 7.10.2.240 (Molecular Devices) in an attempt to separate the contacting cells. Once the applied pressure in the probing pipette is sufficiently high to separate the contacting cells, *F*_*s*_ is calculated from the final pressure and the pressure of the last failed separation using *F*_*s*_= π *R*_*p*_^*2*^ (*P*_*n-1*_+*P*_*n*_)/2, with *R*_*p*_being the micropipette radius, *P*_*n*_the pressure applied by the probing micropipette during the separating step, and *P*_*n-1*_the pressure applied by the probing micropipette during the pulling step preceding the separating step. Doublets that separate during their first pull are not considered since they lack a lower bound to calculate their separation force. The mean duration of separation of the WT doublets included in this study was 33 ± 1 s (mean ± SEM of 85 doublets).

### Data analysis

#### Pipette size, contact size and shape

Doublets are synchronized using the 3^rd^ cleavage division from 4-to 8-cell stage. For the whole embryos obtained previously (44) that are used in this study, the first and last 3^rd^ cleavage divisions occur between 0 and 2 h from each other. To compare the timing of whole embryos and doublets, we used the mean delay between cleavage divisions of whole embryos as the time after 3^rd^ cleavage.

We use FIJI (45) to measure the contact angle of cell-cell contacts in whole embryos and doublets. Using the angle tool, the equatorial plane in which two contacting cells are observed is chosen to manually measure the external contact angle. Data of compacting whole WT embryos are taken from freely available time lapse images obtained previously (44).

To measure the size of contacts on doublets, the line tool of FIJI is used to manually measure length of contact at its equatorial plane.

To measure the size of micropipettes, the line tool of FIJI is used to manually measure the inner diameter near the tip.

#### Immunostaining

Doublets are imaged and synchronized using the 3^rd^ cleavage division from 4-to 8-cell stage before being fixed and stained.

We use FIJI (45) to measure the intensity of Cdh1 and filamentous actin (phalloidin) in doublets to determine their distribution (i) at the cell-cell contact vs the cell-medium interfaces and (ii) at the contact rim vs central disc.

1. To determine the intensity ratio between the contact and cell-medium interface, we measured the intensity I_cc_at cell-cell contacts and intensities I_cm1_and I_cm2_ at the free surface of contacting cells 1 and 2. A 1 μm thick segmented line is manually drawn at the equatorial plane of the cell doublet to measure the mean intensities I_cc_, I_cm1_ and I_cm2_ at the corresponding interfaces.
2. To determine the intensity ratio between the contact rim and central disc, we measured the intensities I_disc_and I_ring_, with the latter one defined as a 1 μm thick circle at the rim of the contact. A 4 μm wide rectangle is drawn along the cell-cell contact to project it along the z axis using 3D project and interpolation functions of FIJI. On the projection, a circle is manually drawn around the entire cell-cell contact to measure its total intensity and area. A 2 μm smaller circle is then positioned onto the center of the cell-cell contact to measure its total intensity and area. To calculate the mean intensity I_disc_, the total intensity of the small circle is divided by its area. To calculate the mean intensity I_ring_, the total intensity of the small circle is subtracted to the one of the large circle, the resulting intensity is divided by the area of the large circle to which the area of the small circle is subtracted.

#### Statistics

Mean, standard deviation, standard error of the mean (SEM), are calculated using Excel (Microsoft). Pearson’s correlation statistical significance is determined on the corresponding table. Statistical significance is considered when *p* < 10^-2^.

The sample size was not predetermined and simply results from the repetition of experiments. No sample was excluded. No randomization method was used. The investigators were not blinded during experiments.

## Results

### Cell doublets as a simplified model for studying mouse embryo compaction

To study compaction in mouse embryos, we simplified the embryo into a minimal adhesive system. As used previously in other contexts (16, 22, 31, 32, 46, 47), cell doublets allow controlling the duration of contact, offer a simple geometry for shape analysis and for understanding the mechanical forces involved in setting the adhesive interface. Importantly, previous studies of dissociated mouse embryos suggested that doublets of mouse blastomeres faithfully and quantitatively recapitulate most aspects of the development of complete embryos such as compaction (6, 48), apicobasal polarization, fate specification, cell internalization (48–51), as well as lumen formation (52, 53).

To quantitatively characterize the compaction of doublets, we dissociated mouse embryos at the 4-cell stage, which, after division to the 8-cell stage, led to the formation of four doublets of 8-cell stage blastomeres per dissociated embryo (Fig 1A). After division, doublets of sister cells formed a cell-cell contact that grew over time, as observed in whole embryos during compaction (Fig 1B, Movie 1, time lapse of whole embryo obtained from (44)). Measuring the external angle of contact of compacting doublets and whole embryos (on images of whole embryos obtained from (44)), we noted similar dynamics and magnitude in the growth of contact angles (Fig 1C), as measured previously (6). Similarly, the radius of contact of doublets increased throughout the 8-cell stage (Fig 1D). In fact, the contact angle and contact radius of doublets follow an identical trend and these were highly correlated (Pearson correlation R = 0.965 for 182 measurements on 41 doublets, p < 10^-2^, Fig 1E), as expected from the geometry of doublets that keep constant volume (6).

**Figure 1:**
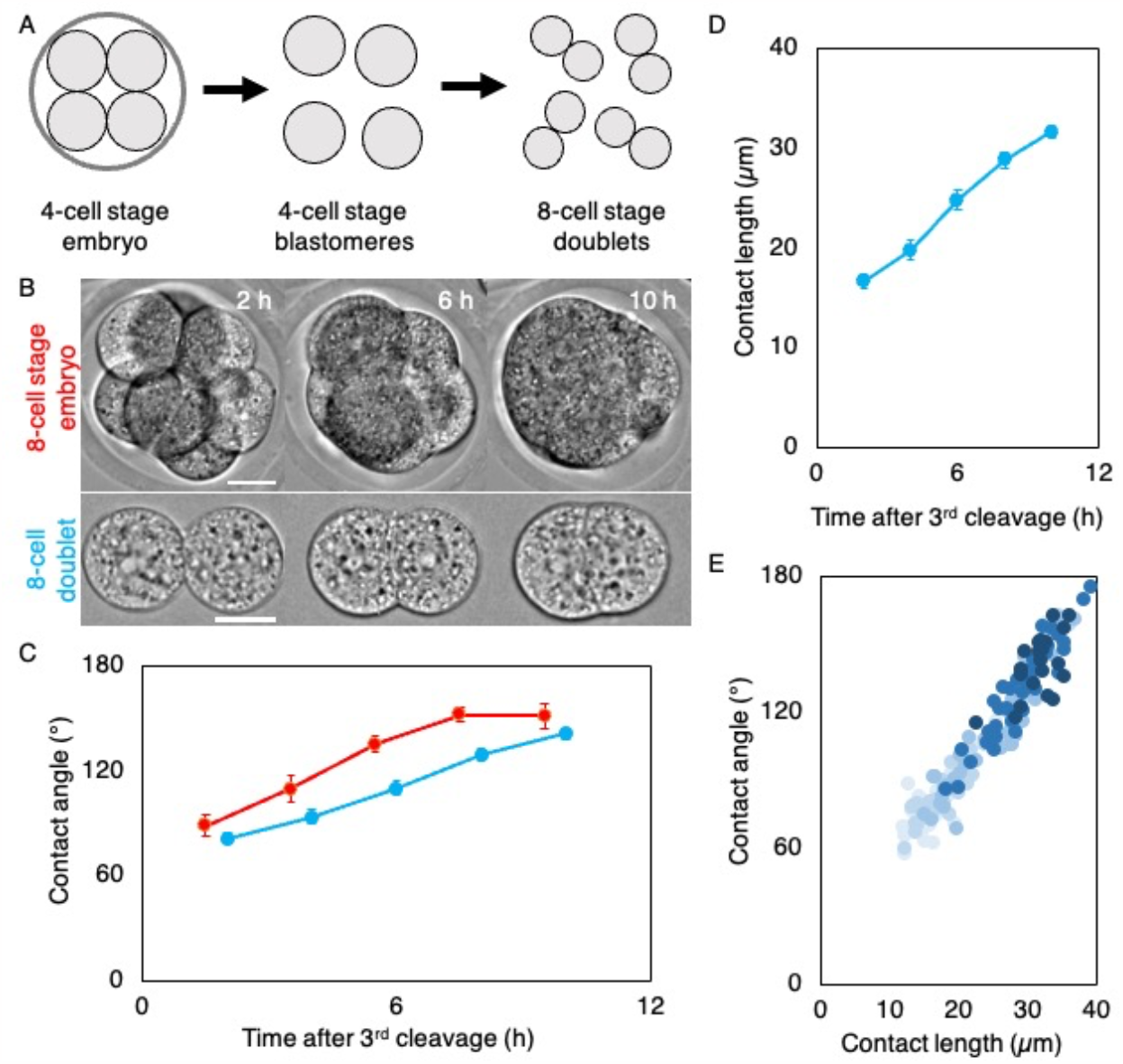
compaction of 8-cell stage doublets. **A)** Schematic diagram of doublet preparation. 4-cell stage mouse embryos are stripped from their zona pellucida and dissociated into individual blastomeres, which then divide into 8-cell stage doublets. **B)** Time lapse images of 8-cell stage embryo and doublet during their compaction (Movie 1). Images of whole embryo from (44). Time after third cleavage is indicated, scale bars are 20 μm. **C)** External contact angle of whole embryos (red) and doublets (blue) during the 8-cell stage. Doublets are synchronized using their 3^rd^ cleavage division. To account for the asynchronous divisions of whole embryos, the time of the last 4-cell stage blastomere to undergo its 3^rd^ cleavage is taken and the mean duration of division to 8-cell stage is added. Mean ± SEM of 20 whole embryos from (44) and of 41 doublets from 11 embryos are shown. **D)** Contact length of 8-cell stage doublet over time after the 3^rd^ cleavage. Mean ± SEM of 41 doublets from 11 embryos are shown. **E)** Contact angle as a function of contact length of 182 measurements throughout the 8-cell stage of 41 doublets from 11 embryos (Pearson correlation R = 0.965, p <10^-2^). Shades of blue indicate the timing of measurement going darker from 2 to 10 h after 3^rd^ cleavage.

Since doublets of 8-cell stage blastomeres quantitatively recapitulate compaction of whole embryos while offering a simpler geometry, we conclude that they constitute an ideal ex vivo system to study physiological changes in cell-cell adhesion.

### Mechanical coupling between cells during compaction

Another advantage of cell doublets is the possibility to separate them to probe the strength of an individual cell-cell contact. To do so, we used a DPA assay, which allows measuring separation forces F_s_ of adhering cells up to hundreds of nanonewtons (31, 32).

Doublets were separated at different times after divisions to measure their separation force F_s_ (Fig 2A, Movie 2). Binning doublets into groups of 2 h after division, we measured that the force required to separate doublets increased from ∼40 nN to ∼70 nN throughout the 8-cell stage (Fig 2C). Therefore, during compaction, cell-cell contacts increase their mechanical stability. As observed previously (Fig 1), the contact angles and radii of doublets increased throughout the 8-cell stage before we separated them (Fig 2B). In fact, contact angles, radii and separation forces F_s_ seem to follow identical dynamics, as confirmed by high correlations between F_s_ and contact length or angle (Pearson correlation R = 0.774 and 0. 737 respectively for 88 doublets, p < 10^-2^, Fig 2D and Fig S1). Therefore, increased contact size could largely explain the increased mechanical stability of cell-cell contacts during compaction.

**Figure 2:**
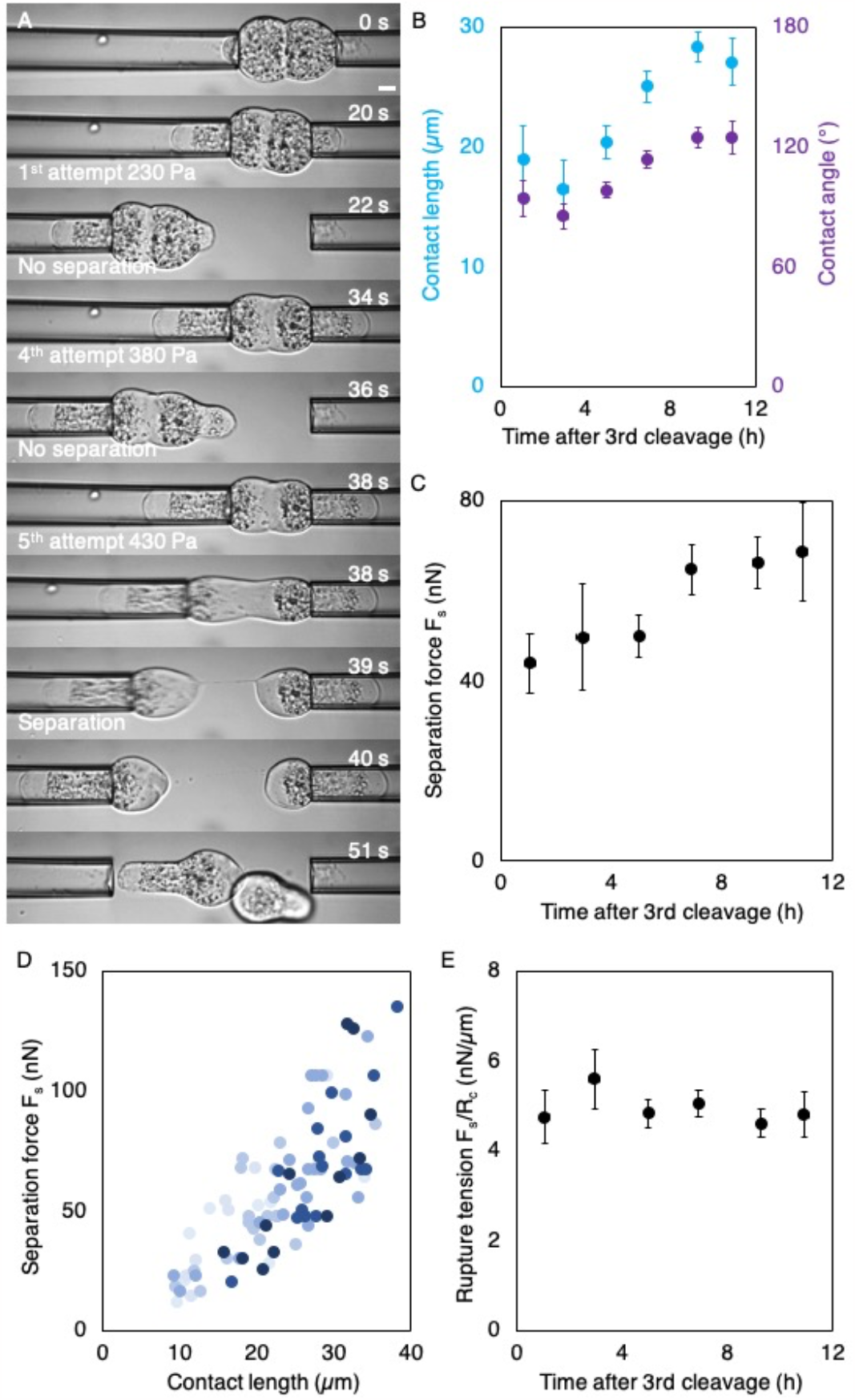
Mechanical coupling of compacting 8-cell stage doublets. **A)** Time lapse images of a doublet separation using dual pipette aspiration (Movie 2). Scale bar, 10 μm. **B)** Contact length (blue) and contact angle (purple) of 8-cell stage doublets just before their separation. Mean ± SEM of doublets in 2 h bins from 0 to 12 h after their 3^rd^ cleavage (n = 9/7/19/25/15/13). **C)** Separation force Fs of 8-cell stage doublets. Mean ± SEM of doublets in 2 h bins from 0 to 12 h after their 3^rd^ cleavage (n = 9/7/19/25/15/13). **D)** Separation force as a function of contact length of 88 measurements throughout the 8-cell stage (Pearson correlation R = 0.774, p <10^-2^). Shades of blue indicate the timing of measurement going darker from 0 to 12 h after 3^rd^ cleavage. **E)** Rupture tension Fs/Rc of 8-cell stage doublets. Mean ± SEM of doublets in 2 h bins from 0 to 12 h after their 3^rd^ cleavage (n = 9/7/19/25/15/13).

### Molecular organization of adherens junctions during compaction

In addition to increased contact size, molecular reorganization of adherens junctions could contribute to the mechanical strengthening of cell-cell contacts. To gain further insights into what explains increased mechanical stability of cell-cell contacts during compaction, we considered different scenarios. In the simplest case, increased contact size allows for more adhesion molecules to engage into adhesive bonds and, with a circular contact with homogeneous adhesive bond density, we would expect mechanical stability to scale with the area of contact. The scaling of mechanical stability with the area of contact may not be linear since in addition to growing in size, cell-cell contacts could recruit and increase the density of adhesion molecules. Furthermore, as measured in other contexts (16, 22), adhesive bonds may become enriched at the rim of cell-cell contacts, where they can more efficiently engage with the actin cytoskeleton and strengthen the mechanical coupling between cells by increasing the effective bond density. In this case, we would expect mechanical stability to scale linearly with the perimeter of cell-cell contacts rather than with the contact area. When comparing the correlations of F_s_ with the perimeter or the surface area of contacts, both yield similarly high correlations (Pearson correlation values R = 0.774 and 0.773 for 88 doublets, respectively, p < 10^-2^), which does not allow us to discriminate between those scenarios. Therefore, we decided to look into the molecular and structural organizations of adherens junctions during compaction.

To investigate the localization of adhesive bonds during compaction, we labelled Cdh1 using immunostaining and filamentous actin using phalloidin on doublets of increasing age (Fig 3A). We measured that actin becomes strongly reduced at cell-cell contacts as compared to the contact-free interface of cells when contacts grow larger during compaction, while Cdh1 shows limited change (Pearson correlation values R = -0.742 and 0.329 for 45 doublets, p < 10^-2^and > 10^-2^between contact angles and actin or Cdh1, respectively, Fig 3B-C, S2A, S2C). This is similar to previous measurements on mouse and human whole embryos (6, 19). Furthermore, we measured a progressive enrichment of Cdh1, but not of actin, at the periphery as compared to the inside of the contact disc when contacts grow larger during compaction (Pearson correlation values R = 0.266 and 0.496 for 45 doublets, p > 10^-2^and < 10^-2^between contact angles and actin or Cdh1, respectively Fig 3D, S2B, S2D), as observed in other doublets (16, 22). Therefore, while Cdh1 does not seem to be further recruited to cell-cell contacts during their expansion, adhesion complexes reorganize to form a ring at the contact edges where the remaining actin cytoskeleton is found. We conclude that, in addition to contact growth, increased mechanical stability may be associated with the accumulation of adhesive bonds at the contact rim.

**Figure 3:**
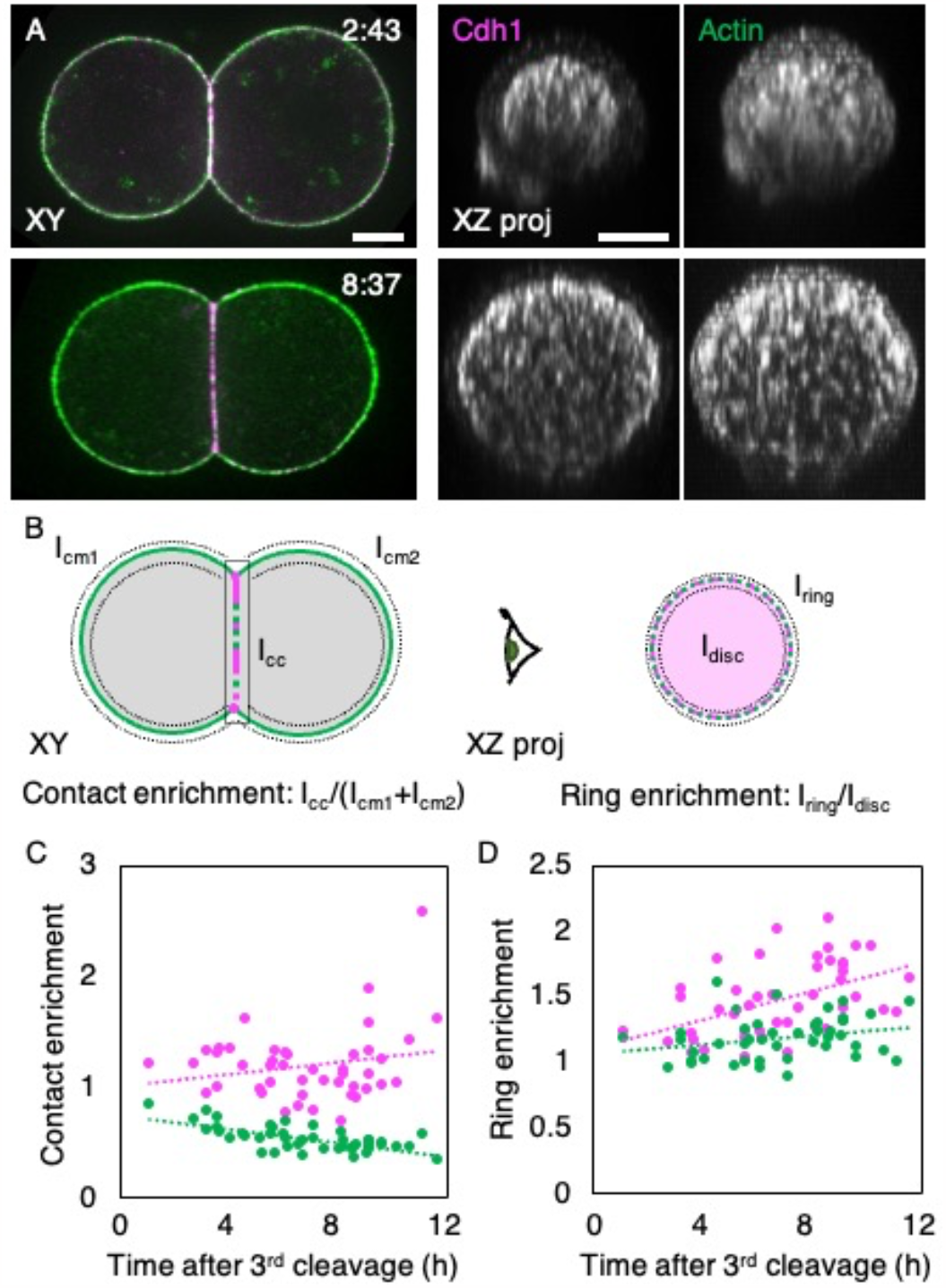
Structural organization of adherens junctions of 8-cell stage doublets. **A)** Immunostaining of early (top) and late (bottom) 8-cell stage doublets. Left shows the merged signals of Cdh1 (magenta) and Actin (green) seen through the equatorial confocal slice of doublets and right shows an orthogonal 4 μm projection of individual signals. Time after third cleavage is indicated, scale bars are 10 μm. **B)** Schematic diagram of quantifications characterizing the molecular organization of doublets: contact (left) and ring (right) enrichments. Left shows the equatorial plane in which intensities are measured along the cell medium interfaces of both cells Icm1 and Icm2 as well as the intensity along the contact Icc using a 1 μm thick segmented line. Contact enrichment is calculated as Icc / (Icm1 + Icm2). Right shows an en face view of the cell-cell contact on which the intensities of a 1 μm peripheral ring Iring and of the inner disc Idisc are measured. Ring enrichment is calculated as Iring / Idisc. **C-D)** Contact (C) and ring (D) enrichments of Cdh1 (magenta) and Actin (green) throughout the 8-cell stage of 45 doublets (Pearson correlations R = 0.216 (p > 10^-2^) and 0.475 (p < 10^-2^) for Cdh1; -0.680 (p < 10^-2^) and 0.266 (p > 10^-2^) for Actin).

To test the potential influence of the structural reorganization of adhesion complexes during compaction without the contribution of contact expansion, we normalized the separation force to the radius of contact R_c_. This gives a rupture tension F_s_/R_c_, which may reflect an effective bond density (16). We observed no change in rupture tension F_s_/R_c_ during compaction (Fig 2E), indicating that the enrichment of Cdh1 at the contact rim is not associated with stronger effective bond density during mouse embryo compaction. Instead, increased mechanical stability during compaction is primarily explained by increased contact size.

### Reduced mechanical coupling in embryos lacking Cdh1-mediated adhesion

Since Cdh1 mediates cell-cell adhesion during compaction, we reasoned that reducing its levels, should reduce the mechanical stability of cell-cell contacts. Maternal heterozygous Cdh1 mutant (mCdh1^+/-^) embryos should have less Cdh1 at the time of compaction since they lack maternal Cdh1 and can only rely on zygotic expression from their paternal allele. Lower levels of Cdh1 should likely decrease the mechanical stability cell-cell contacts and potentially their effective bond density. Moreover, compaction is impaired in mCdh1^+/-^ embryos, which form smaller cell-cell contacts than WT embryos (6). With both smaller contacts and fewer adhesion molecules, the effective bond density of mCdh1^+/-^ embryos is difficult to predict.

To characterize this, we generated doublets from mCdh1^+/-^ embryos (Fig 4A, Movie 3). Binning doublets over groups of 4 h after division, we found that mCdh1^+/-^ doublets start with smaller contacts than WT and compact less (Fig 4B), as observed previously in whole embryos (6). As expected from the smaller contacts, we also measured lower separation forces F_s_ in mCdh1^+/-^ doublets than in WT ones (Fig 4C). Nevertheless, mCdh1^+/-^ doublets could grow from ∼30 to ∼40 nN during the 8-cell stage (Fig 4C). Therefore, mCdh1^+/-^ doublets reach, at the end of their 8-cell stage, a level of mechanical coupling comparable to WT doublets at the start of their 8-cell stage (Fig 4D). This could be explained by mCdh1^+/-^ doublets catching up to contact sizes of WT doublets. However, when measuring the rupture tension F_s_/R_c_ for mCdh1^+/-^ doublets at the beginning of the 8-cell stage, we found that it is higher than in WT (Fig 4E). Then, as mCdh1^+/-^ contacts enlarge and reach the levels of WT at the start of compaction, the rupture tension F_s_/R_c_ decreases to reach WT levels. This suggests that the smallest contacts of mCdh1^+/-^ mutants may have higher effective bond density than contacts formed by WT. Therefore, at low level of expression, the relationship between Cdh1 levels and effective bond density appears complex.

**Figure 4:**
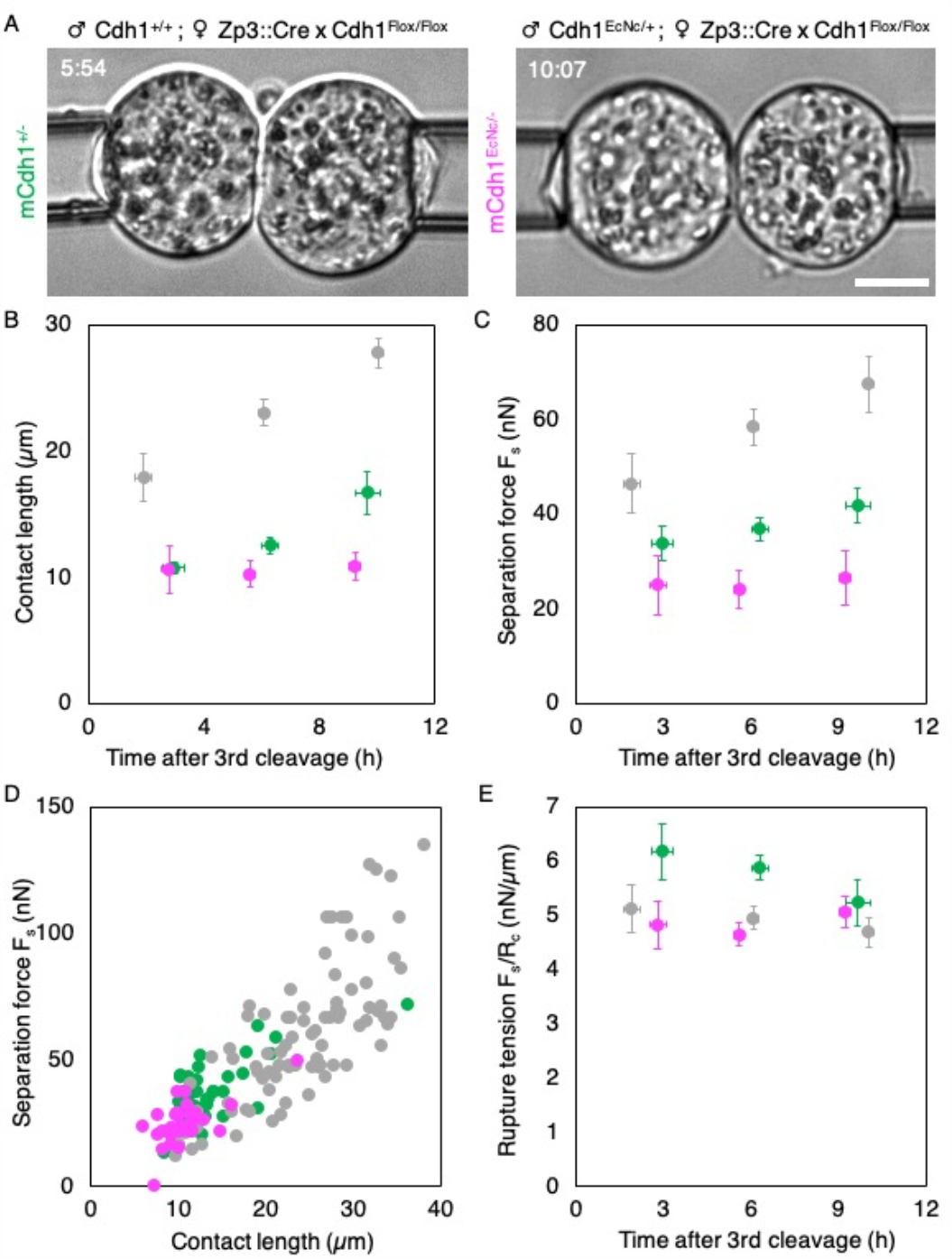
Mechanical coupling of 8-cell stage doublets from mCdh1^+/-^ and mCdh1^EcNc/-^ embryos. **A)** Images of mCdh1^+/-^ (left) and mCdh1^EcNc/-^ (right) 8-cell stage doublets. Time after third cleavage is indicated, scale bar is 10 μm. **B-C)** Contact length (B) and separation force Fs (C) of WT (grey, n = 16/44/28), mCdh1^+/-^ (green, n = 9/16/14) and mCdh1^EcNc/-^ (magenta, n = 9/12/13) 8-cell stage doublets just before their separation. Mean ± SEM of doublets in 4 h bins from 0 to 12 h after their 3^rd^ cleavage. **D)** Separation force as a function of contact length of WT (grey, n = 88), mCdh1^+/-^ (green, n = 39) and mCdh1^EcNc/-^ (magenta, n = 34) doublets throughout the 8-cell stage. **E)** Rupture tension Fs/Rc of WT (grey, n = 16/44/28), mCdh1^+/-^ (green, n = 9/16/14) and mCdh1^EcNc/-^ (magenta, n = 9/12/13) 8-cell stage doublets. Mean ± SEM of doublets in 4 h bins from 0 to 12 h after their 3^rd^ cleavage.

Together, we conclude that reducing the levels of Cdh1 leads to decreased mechanical coupling of cell-cell contacts, which cannot be explained solely by the reduced contact size.

### The intracellular domain of Cdh2 is less effective than that of Cdh1

In addition to the levels of adhesion molecules, their individual mechanical stability is likely to influence the overall mechanical stability of cell-cell contacts. The mechanical stability of adherens junctions is limited by their intracellular coupling rather than the extracellular binding of cadherin adhesion molecules (16, 32). Indeed, as observed in other contexts (16), separating doublets of 8-cell stage blastomeres leads to the formation of long membrane tethers connecting separated cells, indicating that some cadherin adhesion molecules were not separated (Movie 2, Fig 2A). To test the role of intracellular coupling, we took advantage of a transgenic mouse in which the endogenous locus of Cdh1 has been replaced with the cDNA encoding a chimeric protein consisting of the extracellular and transmembrane domains of Cdh1 and the intracellular domain of Cdh2 called EcNc (34). To generate embryos expressing only the chimeric protein at the stage of compaction, we removed Cdh1 maternally and provided the EcNc allele paternally using heterozygous fathers Cdh1^EcNc/+^. To distinguish between mCdh1^+/-^ and mCdh1^EcNc/-^ embryos, we used immunostaining using antibodies targeting either the extra- or intra-cellular domains of Cdh1, or the HA tag added to the EcNc (34).

At the start of the 8-cell stage, the contacts of mCdh1^EcNc/-^ doublets are small, similar in size to the ones of mCdh1^+/-^ doublets (Fig 4A-B, Movie 4). However, unlike mCdh1^+/-^ doublets, mCdh1^EcNc/-^ doublets do not increase their contact size (Fig 4A-B). Therefore, expression of EcNc chimeric cadherin does not elicit compaction, which could be due to reduced signaling or mechanical coupling from the cytoplasmic domain of Cdh2 compared to Cdh1. As expected from their contacts remaining small, mCdh1^EcNc/-^ doublets display overall weak mechanical coupling, which does not increase during the 8-cell stage (Fig 4C). Finally, calculating the rupture tension F_s_/R_c_ in mCdh1^EcNc/-^ doublets revealed that it is lower than in mCdh1^+/-^ doublets at comparable contact size (Fig 4E). This suggests that the intracellular domain of Cdh2 may be less mechanically stable than the one of Cdh1. Together, we conclude that Cdh1 mediated cytoskeletal anchoring is key to elicit compaction and to build mechanically stable cell-cell contacts.

## Discussion

The shaping of the eutherian embryo begins with compaction, a developmentally regulated adhesion process during which cells form a tighter structure. The mechanical characterization of compacting human and mouse embryos previously revealed the changes in surface tensions responsible for contact expansion (15, 54, 6, 19). Here, we studied the mechanical stability of cell-cell contacts throughout compaction using an ex vivo reduced system of cell doublets (Fig 1, Movie 1). We found that, as compaction proceeds, cell-cell contacts require higher forces to be separated (Movie 2). This increased mechanical stability is primarily explained by the spreading of cell-cell contacts associated with compaction (Fig 2). While adherens junctions are remodeled during compaction (Fig 3), this does not seem to be associated with increased mechanical stability since the rupture tension, indicative of effective bond density, does not change (Fig 2E). However, when reducing the levels of adhesion molecules to a level that impairs the expansion of contacts, we measured an effect on contact mechanical stability, which also depends on the molecular nature of the intracellular coupling of adherens junctions (Fig 4). Together, our ex vivo analysis reveals the contributions of contact spreading and molecular reorganization to the mechanical strengthening of cell-cell contacts during compaction.

Compaction is the first morphogenetic movement in eutherian mammals, such as mice and humans (55, 56). In human embryos, weak or delayed compaction is associated with poorer prognosis (57, 58) but could be indicative of other developmental defects such as aneuploidy (59, 60). In fact, mutant mouse embryos, which lack maternal Cdh1 or maternal non-muscle myosin II but express these genes from their paternal alleles (20, 44), fail to compact but form blastocysts and viable mice. Therefore, compaction is not essential to mouse development and its function is unclear. However, defective compaction in some cells of human embryos is associated with their exclusion from the embryo (60), which can eventually cause cells to detach from the compacted embryo, especially in the absence of egg shell (19). Here, we show that contact spreading associated with compaction provides increased mechanical coupling between cells (Fig 2). Therefore, compaction may simply prevent cells from detaching from the embryo.

Surface tension measurements revealed the prominent role of actomyosin contractility in controlling the surface tensions driving compaction in mouse and human embryos (6, 19). Increased contractility and associated surface tensions pulling on adherens junctions have been proposed to lead to their mechanosensitive strengthening in a variety of contexts (22, 25, 26, 61–63). This is associated with the relocation of adhesive bonds to the contact periphery, which forms a ring in doublets (22, 63), as well as in vivo (16). In compacting mouse embryos, for which contractility doubles the surface tensions applied at the edges of cell-cell contacts, we also observe the relocation of Cdh1 in a peripheral ring, without significant recruitment of Cdh1 to cell-cell contact from the remaining of the cell surface (Fig 3). However, this relocation is not associated by increased effective bond density, since the rupture tension F_s_/R_c_ does not change during the formation of an adhesive ring (Fig 2E). We do not observe a significant relocation of the actin cytoskeleton to the adhesive ring (Fig 3), as observed in other systems (16, 22). Since increased actin recruitment is a hallmark of mechanical reinforcement of adherens junction via the mechanosensitive recruitment of junctional proteins such as vinculin (25, 61, 64), the lack of actin recruitment in compacted doublets further indicates the absence of reinforcement of the intracellular coupling of Cdh1 during mouse compaction. Taken together, our experimental results point at a simple explanation for the mechanical reinforcement of cell-cell contact stability during compaction: increased contact size leads to larger peripheral adhesive rings cumulating more adhesive bonds of constant effectiveness.

While effective bond density does not seem to change during compaction, the intracellular coupling of cadherin is essential to their function as a mechanical anchor (16, 32). Different cadherins have distinct mechanical coupling, as measured for example for type I Cdh1 and type II Cdh7 cadherin adhesion molecules using chimeric proteins (33). Here, we looked into the difference between Cdh1 and Cdh2 cytoplasmic domains. A previous study looked into zygotic replacement of Cdh1 with a chimeric protein EcNc made of the extracellular domain of Cdh1 and the intracellular domain of Cdh2, which did not cause compaction defect (34). We eliminated the maternally provided Cdh1, zygotically expressed in its place the chimeric adhesion molecule EcNc and found that 8-cell stage doublets from these embryos do not compact (Fig 4). This indicates that in the initial characterization of zygotic EcNc mutants (34), maternally provided Cdh1 could most likely compensate for the presence of EcNc at the time of compaction. Since zygotically expressed Cdh1 elicits some compaction, the ability of EcNc to function as an adhesion molecule is unclear. EcNc adhesion molecules may fail at signaling to the actomyosin cytoskeleton to allow compact expansion and/or fail at mechanically coupling the cortices of adhering cells. Since we measured weaker rupture tensions when zygotically expressing EcNc than when expressing Cdh1 in maternal Cdh1 mutants (Fig 4), this suggests that EcNc cannot provide much adhesive coupling. Therefore, the intracellular coupling of Cdh2 is likely weaker than the one of Cdh1. Further studies into the specificities of cadherin adhesion molecules signaling and mechanical coupling will be needed to uncouple the distinct functions of cadherins in cell-cell adhesion.

## Acknowledgements

We thank the imaging platform of the Genetics and Developmental Biology Unit at the Institut Curie (PICT-IBiSA@BDD, member of the French National Research Infrastructure France-BioImaging ANR-10-INBS-04) for their outstanding support, in particular Olivier Leroy for his help implementing micromanipulator control into Metamorph; the animal facility of the Institut Curie for their invaluable help. We thank David Rozema for critical reading of the manuscript.

Research in the lab of J.-L.M. is supported by the Institut Curie, the Centre National de la Recherche Scientifique (CNRS), the Institut National de la Santé Et de la Recherche Médicale (INSERM), and is funded by grants from the European Research Council Starting Grant ERC-2017-StG 757557, the European Molecular Biology Organization Young Investigator program (EMBO YIP), the INSERM transversal program Human Development Cell Atlas (HuDeCA), Paris Sciences Lettres (PSL) QLife (17-CONV-0005) grant and Labex DEEP (ANR-11-LABX-0044) which are part of the IDEX PSL (ANR-10-IDEX-0001-02).

## Author contributions

L.P. performed experiments and prepared data for analyses. L.P. and J.-L.M. designed the project, analyzed the data and wrote the manuscript. J.F. helped with data analysis. J.-L.M. acquired funding.

## Supplementary figures

**Supplementary figure 1:**
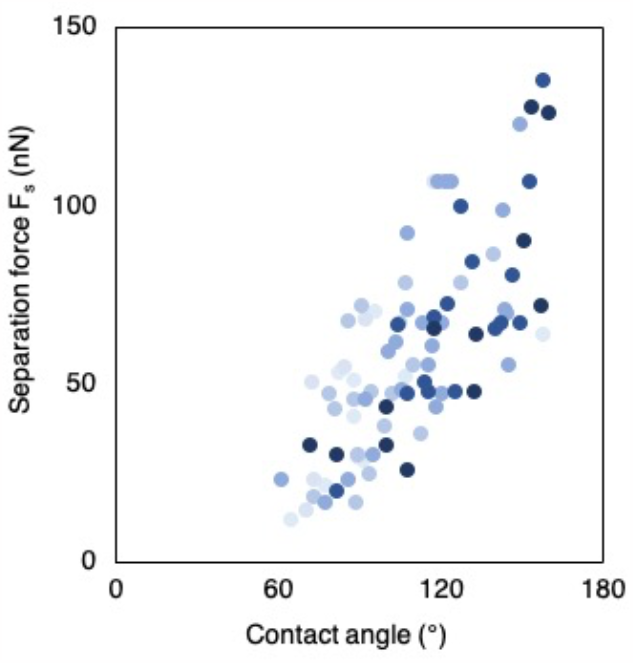
Relationship between separation force and contact angle of 8-cell stage doublets. Separation force as a function of contact angle of 88 measurements throughout the 8-cell stage (Pearson correlation R = 0.737, p <10^-2^). Shades of blue indicate the timing of measurement going darker from 0 to 12 h after 3^rd^ cleavage.

**Supplementary figure 2:**
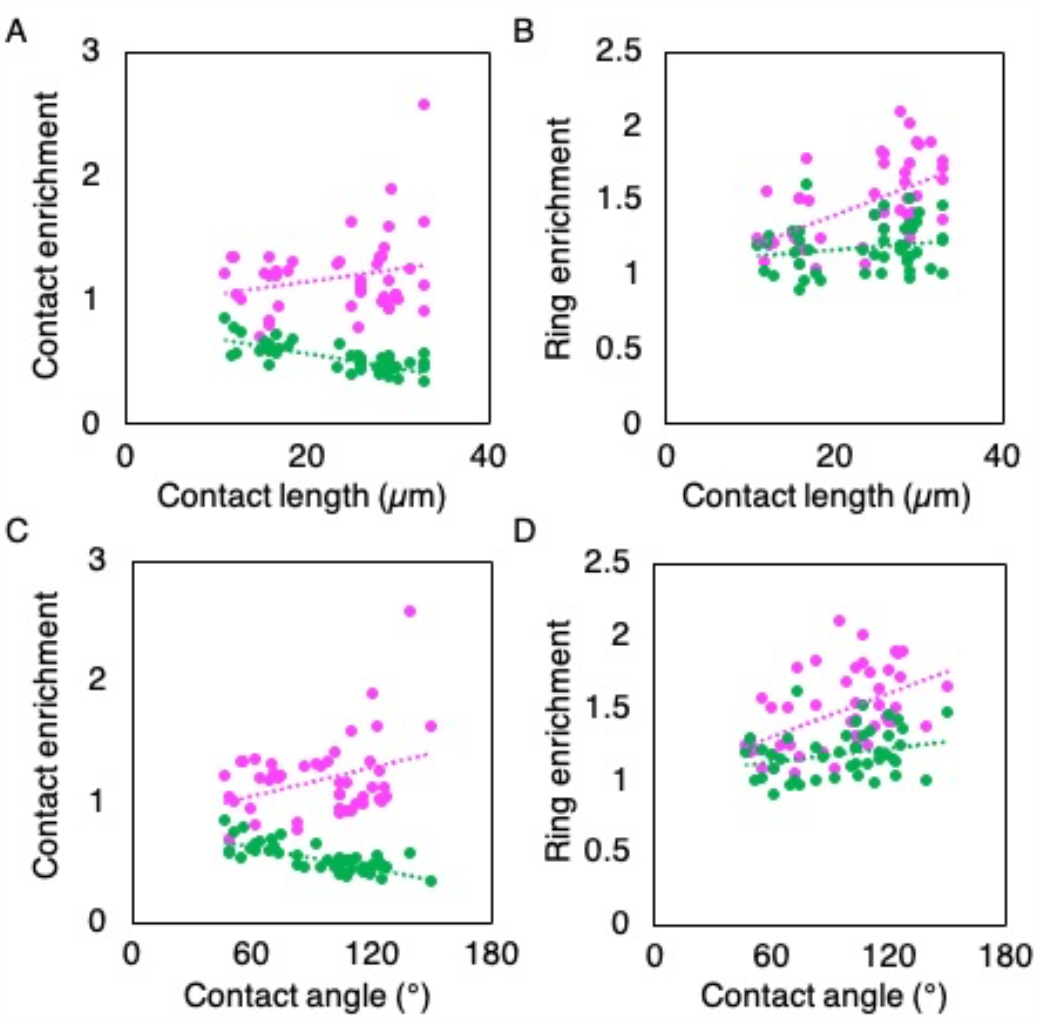
Structural organization of adherens junctions of growing doublets. **A-D)** Contact (A, C) and ring (B, D) enrichments of Cdh1 (magenta) and Actin (green) as a function of contact length (A, B) and angle (C, D) of 45 doublets (Pearson correlations (A) R = 0.223 (p > 10^-2^) for Cdh1 and - 0.747 (p < 10^-2^) for Actin; (B) R = 0.543 (p < 10^-2^) for Cdh1 and 0.196 (p > 10^-2^) for Actin; (C) R = 0.329 (p > 10^-2^) for Cdh1 and -0.742 (p < 10^-2^) for Actin; (D) R = 0.496 (p < 10^-2^) for Cdh1 and 0.266 (p > 10^-2^) for Actin.

## Movie legends

**Movie 1: Time lapse of a compacting whole embryo and of a doublet**. Images of a whole embryo (left) and of a doublet (right) undergoing their 3^rd^ cleavage division and compacting. Images are acquired every 30 min, scale bar 20 μm.

**Movie 2: Time lapse of separation of a WT doublet using dual pipette aspiration**. Images of a WT doublet being separated using two pipettes. The left pipette is set to 400 Pa and is moved to the left at 60 μm/s after the right pipette is placed in contact with the right hand side cell of the doublet. The pressure of the right pipette is increased to 230, 280, 330, 380 and finally 430 Pa between pulls. Images are acquired every 100 ms, scale bar 10 μm.

**Movie 3: Time lapse of separation of a mCdh1^+/-^ doublet using dual pipette aspiration**. Images of a mCdh1^+/-^ doublet being separated using two pipettes. The right pipette is set to 210 Pa and is moved to the right at 60 μm/s after the left pipette is placed in contact with the left hand side cell of the doublet. The pressure of the left pipette is increased to 60, 110, 150, and finally 200 Pa between pulls. Images are acquired every 100 ms, scale bar 10 μm.

**Movie 4: Time lapse of separation of a mCdh1^EcNc/-^ doublet using dual pipette aspiration**. Images of a mCdh1^EcNc/-^ doublet being separated using two pipettes. The right pipette is set to 245 Pa. The left pipette is moved to the left at 60 μm/s after the right pipette is placed in contact with the right hand side cell of the doublet. The pressure of the left pipette is increased to 100, 150, 200 and finally 250 Pa between pulls. Images are acquired every 100 ms, scale bar 10 μm.

